# The dominance of non-native plants over native plants increases with the number of global change factors

**DOI:** 10.1101/2024.12.22.630009

**Authors:** Xiong Shi, Duo Chen, Mark van Kleunen

## Abstract

Global environmental change and plant invasion are both recognized as key indicators of the Anthropocene, with the former potentially accelerating the latter. Still, whether multiple co-acting global change factors (GCFs) influence plant invasions even stronger remains untested despite the fact that in nature GCFs usually act together rather than alone. Here, four non-native and four native species were cultivated jointly as communities in mesocosm pots, and we tested their responses to increasing GCF numbers (0, 1, 2, 3, 4, 5). We found that GCF number had no influence on community productivity but increased the biomass proportion of non-native plants. GCF number increased biomass but had no influence on height for non-native plants, whereas both biomass and height of native plants decreased with increasing GCF number. Consequently, our findings indicate that dominance may increases when habitats experience multiple global change factors, and thus that invasion risk of non-native plants could potentially be underestimated from studies on single GCFs.

**Open Research statement:** The data supporting the results will be made available upon acceptance.

## 1. Introduction

The Anthropocene is characterized by various global environmental changes, including pollution by microplastics, the presence of artificial light at night (ALAN), increased nitrogen deposition, the frequency of drought events, and so forth (Lewis & Maslin, 2015). It has been shown that many of these global change factors (GCFs) can have considerable effects on plant growth and plant communities (Choat *et al*., 2018; Rillig *et al*., 2019a; Ma *et al*., 2020; Feeley *et al*., 2020; Speißer *et al*., 2021). For example, ALAN can promote plant invasion (Liu *et al*., 2022) and decrease community diversity and productivity (Bucher *et al*., 2023), nitrogen deposition can increase photosynthetic carbon gain and decrease water loss (Liang *et al*., 2020), and drought can result in tree mortality (Choat *et al*., 2018) and reduce vegetation productivity (Xu *et al*. 2019).

However, in nature, these GCF factors usually do not act in isolation but many of them act simultaneously, and their combined effects may be more complex, possibly resulting in synergistic and antagonistic effects (Rillig *et al*., 2019b; Zandalinas *et al*., 2021; Speißer *et al*., 2022). Therefore, one of the major current topics in ecology is understanding how multiple simultaneously acting GCFs affect the performance of individual plant species and plant communities.

Consequently, in recent years, there has been a growing interest in how ecological processes are affected by the number of multiple GCFs acting together (Rillig *et al*., 2019b; Sáez-Sandino *et al*., 2024; Zhou *et al*., 2024). For instance, Zandalinas *et al*. (2021) found that multiple GCFs caused a more critical decline in plant growth and survival than single GCFs did. Similarly, another recent study found that simultaneous GCFs reduced the diversity and evenness of native mesocosm communities (Speißer *et al*., 2022). However, previous studies have focused on how native plants or native communities are affected by multiple GCFs. In fact, human activities have not only resulted in various environmental changes, but have also caused an increasing number of plant species to establish persistent populations outside their native ranges (van Kleunen *et al*., 2015). Some of these non-native plant species may even become invasive and dominate native communities, consequently resulting in a reduction of native community diversity (Vilà *et al*., 2011; Carboni *et al*., 2021). Understanding the effects of the number of multiple GCFs on invaded communities is crucial to predict non-native plant invasion under complex global change scenarios.

It has been suggested that non-native plants often perform better than native plants under various environmental conditions, as they have a so-called Jack-and-Master strategy (Richards *et al*., 2006; Molina-Montenegro *et al*., 2018). For single environmental factors, many non-native plants are also more responsive than native ones (Liu *et al*., 2017, 2022; Kimball *et al*., 2024).

As a consequence, the likelihood that native communities will be invaded by non-native species (i.e. community invasibility) is likely to increase with the simultaneous action of multiple GCFs. However, still little is known about how non-native and native plants growing together respond to increasing numbers of GCFs.

We hypothesize that plant communities become more susceptible to non-native plant invasions with increasing GCF numbers (Fig 1A). There are five potential scenarios that could underlie such a GCF-number effect (Fig 1B): (I) performance of non-native plants increases while that of native plants decreases; (II) performance of non-native plants increases while that of native plants changes less; (III) performance of non-native plants changes little while that of native plants decreases more; (IV) performance of both increases but more strongly for non-native plants than for native ones; (V) performance of both decreases but less strongly for non-native plants than for native ones.

**Figure 1.**
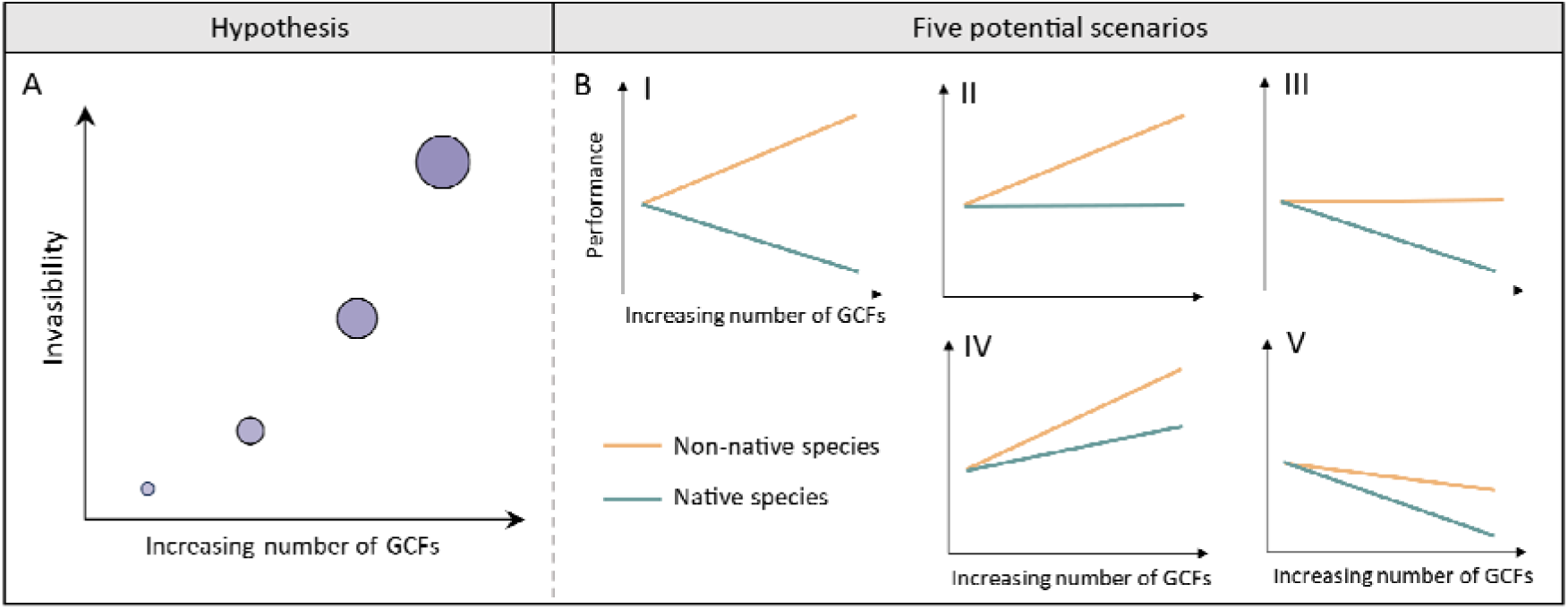
Illustration of the hypothesized positive effect of the number of simultaneously acting global change factors (GCFs) on invasibility or on the dominance of non-native species in already invaded communities (A), and five possible performance scenarios of non-native and native that underly the hypothesized relationship (B): I, performance of non-native plants increases while that of native plants decreases; II, performance of non-native plants increases but that of native plants changes less; III, performance of non-native plants does not change but that of the native plants decreases; IV, performance increases but more strongly for non-native plants than for native plants; V, performance decreases but less strongly for non-native plants than for native plants.

To test the effects of multiple GCFs on non-native and native plants, we conducted a greenhouse experiment by constructing a community containing four non-native and four native plant species. Thereafter, these mesocosm communities were exposed to five single-GCF treatments (ALAN, drought, microplastic pollution, nitrogen deposition and salinization) or treatments with different GCF numbers (levels 2, 3, 4 and 5). We addressed the following main questions: (1) How does an increase in GCF number affect community productivity and the relative dominance of non-native plants? (2) Which of the scenarios we describe in Fig. 1 is more closely aligned with the actual responses of non-native and native plants to GCF numbers? (3) Are there species-specific responses among non-native and native species to the GCF number? (4) Do some of the five single factors have specific effects? If so, to what extent do such factors contribute to the effect of GCF number?

## 2. Material and methods

### 2.1 Study species

To create a community, we selected eight co-occurring plant species, of which four are native to Germany (*Bromus sterilis* L., *Daucus carota* L., *Plantago lanceolata* L., *Silene vulgaris* (Moench) Garcke) and four are non-natives (*Amaranthus retroflexus* L., *Lepidium virginicum* L., *Lolium multiflorum* Lam., *Senecio inaequidens* DC.), the standards of whether they are native or non-native are based on POWO database (POWO, 2024). To increase the generality of our results, the eight plant species represent seven families and were selected from three life history groups (annual, biennial and perennial; details can be found in Table S1).

### 2.2 Experimental set-up

The experiment was conducted in the greenhouse of the Botanical Garden at the University of Konstanz, Germany (47°69’N, 9°17’E). On 21 August 2023, the seeds of eight species were sown into trays (18 cm × 14 cm × 5 cm) filled with potting soil (Topferde, Einheitserde Co). The trays were placed in a greenhouse with a temperature maintained between 18°C and 25°C. For the soil substrate, we filled each of 96 pots (5 L, 18 cm × 18 cm × 18 cm) with 4 kg of a 1:1 mixture of potting soil and sand. Two weeks after sowing, on 4 September 2023, we transplanted one seedling of each of the eight species into the pots. To avoid potential effects of seedling positions within the pot on their subsequent growth, the four non-native and the four native plants were randomly assigned to planting positions in the pots (Fig S1A). Seedlings that died during the first week after transplanting were replaced. Thereafter, the global change treatments were applied.

#### Global change treatments

We imposed five global change factors (Fig. S1D): drought, salinization, nitrogen deposition, microplastic pollutions and artificial light at night (ALAN). These factors were selected due to their prevalence and the likelihood of their continued increase in the future (Galloway *et al*., 2004; Geyer *et al*., 2017; Kyba *et al*., 2017; Hassani *et al*., 2021; IPCC, 2023). Although there were also other global change factors, such as warming, we did not include them due to the unavailability of facilities at the time. For the drought treatment, we provided only 200 ml of water twice a week to each of the pots, whereas we provided 400 ml of water twice a week to each of the drought-free pots, as in Haeuser *et al*. (2019). The soil-moisture contents of all pots were measured every week (Fig. S2). For the pots exposed to the salinization treatment, we added a 0.5% NaCl solution to the pot every week. We estimated soil salinity by measuring the soil electrical conductivity, and kept the electrical conductivity at a value of approximately 6 ds m^-1^ for the salinization-treated pots (Richards, 1954; Speißer *et al*., 2022). To ensure consistent conductivity across the drought and well-watered treatments, we applied 200 ml of the NaCl solution to drought-stressed pots and 400 ml of the NaCl solution to the well-watered pots, and measured soil electrical conductivity weekly (Fig. S3). Furthermore, the nitrogen deposition treatment was done by mixing into the substrate of the corresponding pots 0.523 g of slow-release nitrogen fertilizer (Floranid® N 31 31-0-0) prior to planting the seedlings. This quantity was calculated based on the average annual nitrogen deposition in nature (50 kg ha^-1^ yr^-1^; Galloway et al. 2004). For the pots containing microplastics, we had mixed 70.72 g of granules (1.0-2.5 mm in diameter) of ethylene propylene diene monomer (EPDM) into the soil substrate of the corresponding pots prior to transplanting the seedlings, resulting in a volume concentration of approximately 2% (Speißer *et al*., 2022; Speißer & van Kleunen, 2023). For the ALAN treatment, we used LED light strips and set the light intensity to 24.5-25 lux between 9 pm and 5 am each day to simulate summer conditions under street lights. To prevent the ALAN-free pots from being disturbed by nearby LED light, wooden boxes (240 cm × 28 cm × 60 cm) that were open at the top and bottom were constructed and placed around the ALAN pots (Fig. S1B). Eight ALAN pots were placed in each of the six ALAN boxes. To account for the potential effect of the wooden boxes on plant growth, the ALAN-free pots were also surrounded by such boxes but without the LED light. Thus, there were six blocks in total, each of which consisted of one ALAN box and one ALAN-free box (Fig. S1C).

#### Combinations of simultaneously acting GCFs

To test the effects of GCF numbers on non-native and native plants, we used six GCF-number levels (0, 1, 2, 3, 4 and 5 GCF; Fig. S1C). For the level of zero-GCF, i.e. the control treatment, no GCF was applied. For levels one, two, three and four, there were either five single GCF treatments (GCF level one) or five GCF combinations for each of the GCF levels two, three and four. For GCF level five, all five GCFs were combined, thus there was only one combination. We created the GCF combinations randomly with the restriction that each GCF was used an equal number of times in each GCF-number level (Fig. S1C). In each block, each GCF-number level was present, and there was a maximum of one replicate per GCF combination. For each treatment combination, we had in total across all blocks at least four replicates. However, to use all the available space and keep the pot number consistent in each box—since some treatments involved only one combination in ALAN or ALAN-free conditions—we had six replicates for the following four treatments: control, ALAN, Drought + Salinization + Nitrogen + Microplastic, and Drought + Salinization + Nitrogen + Microplastic + ALAN, resulting in 96 pots in total (Fig. S1D). All pots were individually placed on plastic dishes (Ø = 26 cm) and were re-randomized within each box after five weeks of growth.

### 2.3 Measurements

To investigate the responses of non-native and native plants to single and multiple GCFs, we used height and biomass as indicators of plant performance (Younginger *et al*., 2017; Xiao *et al*., 2021; Quan *et al*., 2024). On 7 November 2023, two months after the start of the experiment, we observed that the surrounding species are closely intertwined and interacting with one another in the pots, and then we measured the height of each plant individually (from the base to the tip of the shoot) and harvested the above- and belowground biomass of each plant separately. The plant materials were dried at 70 over 72 hours and then weighed.

### 2.4 Statistical analysis

All analyses were conducted in R 4.3.2 (R Core Team, 2024). To test the effects of GCF number on community productivity and biomass proportion of the non-native plants, we fitted two linear mixed models using the *lme* function in the ‘nlme’ package (Pinheiro *et al*., 2024). Total biomass per pot and proportional biomass of non-native plants were included as the response variables, respectively. In these two models, GCF number was used as a fixed continuous factor. To account for variation in plant growth among boxes and among the combinations of GCFs, the experimental block and the identity of the GCF combinations (including the single GCF treatments) were included as random factors. Similarly, to test the effects of the five single GCFs on biomass per pot and proportional biomass of non-native plants, we also ran two models using the subset of pots with the control and the five single GCF treatments. We created five dummy contrasts to split up the difference between control and each of the five single GCF treatments (Schielzeth, 2010). The five dummy contrasts were used as the fixed factor, and the experimental block was treated as a random factor. To improve normality of the residuals, we used a log transformation for the proportional biomass of non-native plants. The significance of fixed effects was assessed using the *Anova* function (likelihood-ratio tests, type II) of the ‘*car’* package (Fox & Weisberg, 2018).

Furthermore, we fitted two models to test whether the responses to GCF number differed between non-native and native plant individuals. Plant height and natural-log-transformed total biomass were used as response variables, respectively. We included species origin (non-native or native), GCF number and their interaction as fixed factors, and included species identity, pot identity, block and the identity of the GCF combination as random factors. In addition, to test whether the effects of various single GCFs on non-native and native plants differed, we further fitted two models using the subset of pots with the control and the five single GCF treatments.

Plant height and square-root-transformed total biomass were used as response variables. Species origin, five dummy contrasts and their interactions were included as fixed factors. Species identity, block and pot identity were included as random factors.

To test the effects of GCF number on each specific study species, we also ran models using a subset of the data for each of the eight species separately. In these models, either plant height or total biomass was included as a response variable. To improve normality of the residuals, we did different transformations (details can be found in Table S2). GCF number was used as a fixed continuous term, and the identities of GCF combinations and blocks were used as random terms.

In addition, as the pattern of GCF-number effects on plant performance may potentially be influenced by the selection effect (i.e. sampling effect) of the specific GCF, as in studies on biodiversity ecosystem functioning (Loreau & Hector, 2001), we assessed this possibility by comparing models with and without the GCF combinations of the interest. Specifically, if we found significant effects of one or more single GCFs on plant performance at both the community and individual levels, we would remove the GCF combinations including the potentially influential factor from the dataset, and then re-run the model to test whether the results remained qualitatively the same. Moreover, to test whether significant GCF-number effects were determined by GCF identities or the interactions between GCFs (i.e. whether the effects of GCFs are additive or non-additive), we also ran hierarchical models to compare which model was the best. This was done based on the hierarchical diversity-interaction modelling framework (Kirwan *et al*., 2009), and details about the model comparisons can be found in Fig. S4.

## 3. Results

### 3.1 Community productivity and biomass proportion of non-native plants

Analysis of the subset of single GCFs revealed that GCFs had no significant influence on community productivity (Table S3, Fig. S5A), while the presence of microplastics tended to increase the biomass proportion of non-native plants (*p* = 0.067, Table S3, Fig. S5B). Analysis of the full dataset showed that an increase in the number of GCFs had no significant influence on the total biomass production of the entire community (Table S4, Fig. 2A). However, the biomass proportion of non-native plants increased with the number of simultaneously acting GCFs (*p* = 0.002, Table S4, Fig. 2B). Hierarchical diversity-interaction analysis for biomass proportion of non-native plants revealed that the GCF identity model fitted the data best (Table 1). The positive GCF-number effect, however, was still significant after removal of the most influential GCF factor (i.e. microplastics) from the model (*p* = 0.024, Table S5, Fig. S6). These results thus indicate that the positive effects of the GCF number on the biomass proportion of non-native plants could be attributed to additive effect of all GCFs.

**Figure 2.**
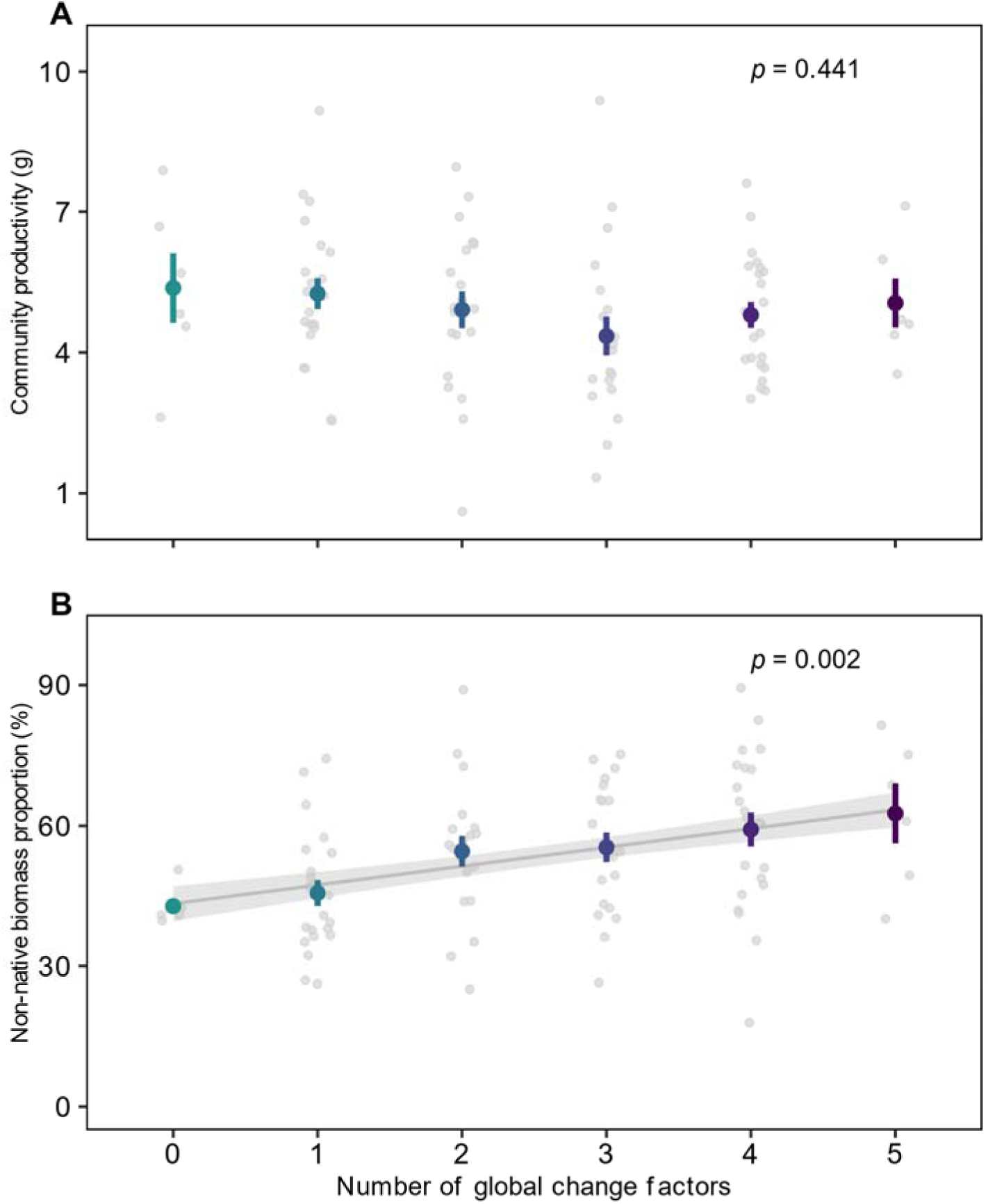
GCF-number effects on total community productivity (A) and on biomass proportion of non-native plants (B). Data are presented as mean values ± 1 SE. The smaller light purple points indicate raw data distributions. In B, the line indicates the significant relationship with its 95% confidence interval. The *p* values indicate the significance of the GCF-number effects.

**Table 1.** Contributions of GCF identity and GCF interactions to GCF-number effects on the biomass proportion of non-native plants (see Figure S4 for an explanation of the different models). Model comparisons were based on the log-likelihood-ratio test, and the number in brackets after each model indicates the reference model. The best-fitting model (i.e. with the lowest AIC) is highlighted in bold.

| Model (reference model number) | AIC | logLik | L.ratio | *P* value |
| --- | --- | --- | --- | --- |
| 1 Null | 770.089 | -381.044 | - | - |
| **2 GCF identity (1)** | **745.698** | **-363.849** | **34.391** | **< 0.001** |
| 3 Separate-pairwise interaction (4) | 752.809 | -357.404 | 12.316 | 0.196 |
| 4 Average GCF interaction (2) | 747.125 | -363.563 | 0.573 | 0.449 |
| 5 Additive GCF-specific interaction contributions (3) | 746.585 | -359.293 | 3.776 | 0.582 |

### 3.2 Growth responses of non-native and native plants to multiple GCFs

Among the single GCF effects on plants, salinization decreased plant height (*p* = 0.006) and tended to decrease total biomass (*p* = 0.088), irrespective of plant origins (Fig. S7, Table S6). The application of artificial light at night tended to promote the height of native plants but to decrease that of non-native plants (*p* = 0.053, Table S6, Fig. S7B). Furthermore, non-native and native plants showed different responses to an increase in GCF number. Specifically, the biomass of non-native plants increased as the GCF number increased, whereas the opposite was true for native plants (*p* = 0.002, Table S7, Fig. 3A). Plant height decreased with an increase in GCF number, but this decrease was stronger for native plants than for non-native plants (*p* = 0.005, Table S7, Fig. 3B). The hierarchical diversity-interaction models showed that for both the biomass and height of non-native plants, the GCF identity model had the best fit (i.e. lowest AIC; Table 2). However, we did not observe significant positive effects of any of the single GCFs on performance of non-native plants (Fig. S7), indicating that only the combined effect of multiple GCFs affected the performance of non-native plants. On the other hand, for biomass and height of native plants, the average GCF interaction model and the separate-pairwise interaction models, respectively, had the best fit (Table 2), suggesting that interactions in general or specific GCF interactions caused non-additive effects for native plant performance.

**Figure 3.**
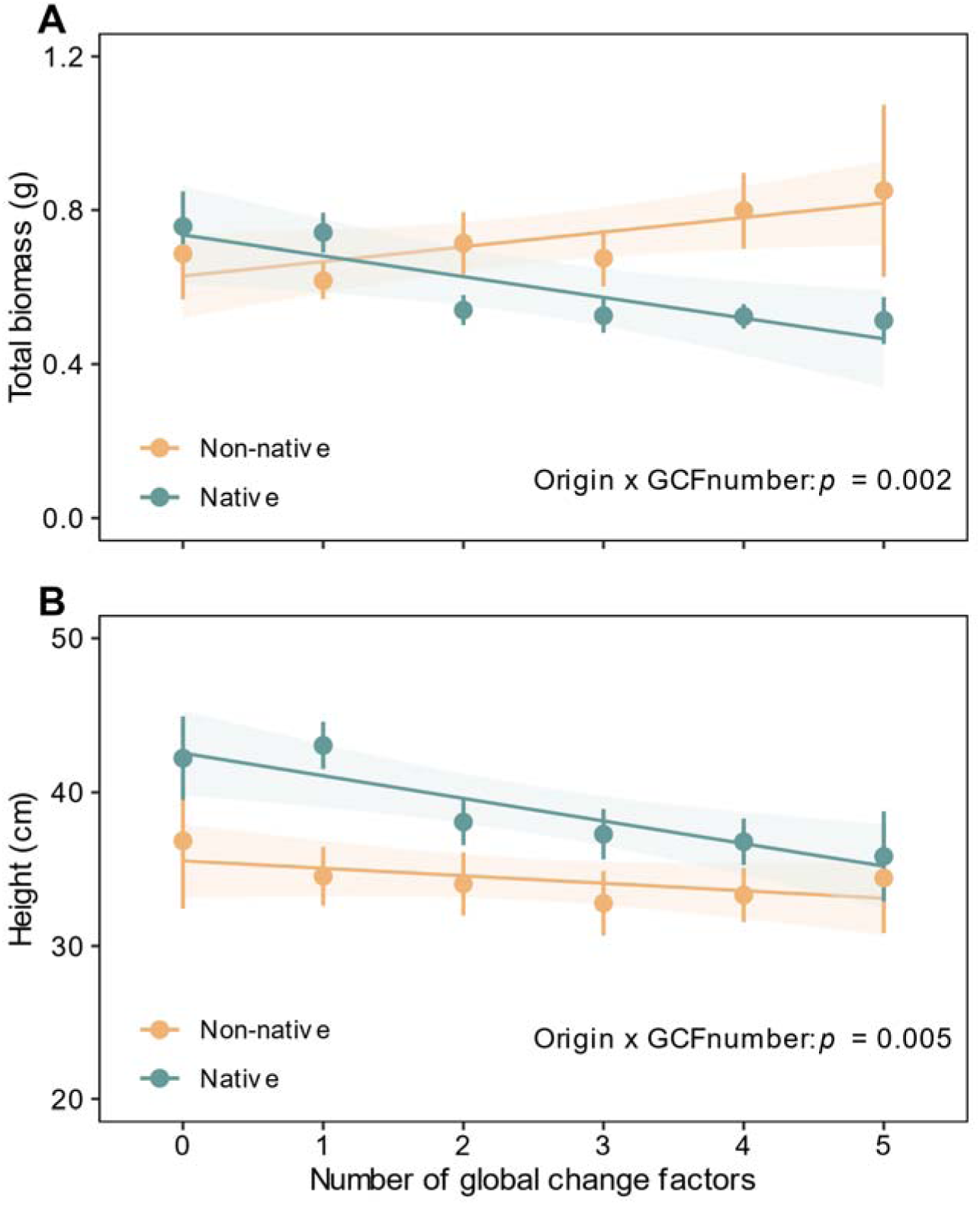
GCF-number effects on total biomass (A) and height (B) of non-native and native plants. Data are presented as mean values ± 1 SE. The lines indicate the fitted relationships for non-native and native plants with their 95% confidence intervals. The *p* values indicate the significance of the Origin × GCF-number interaction.

**Table 2.**
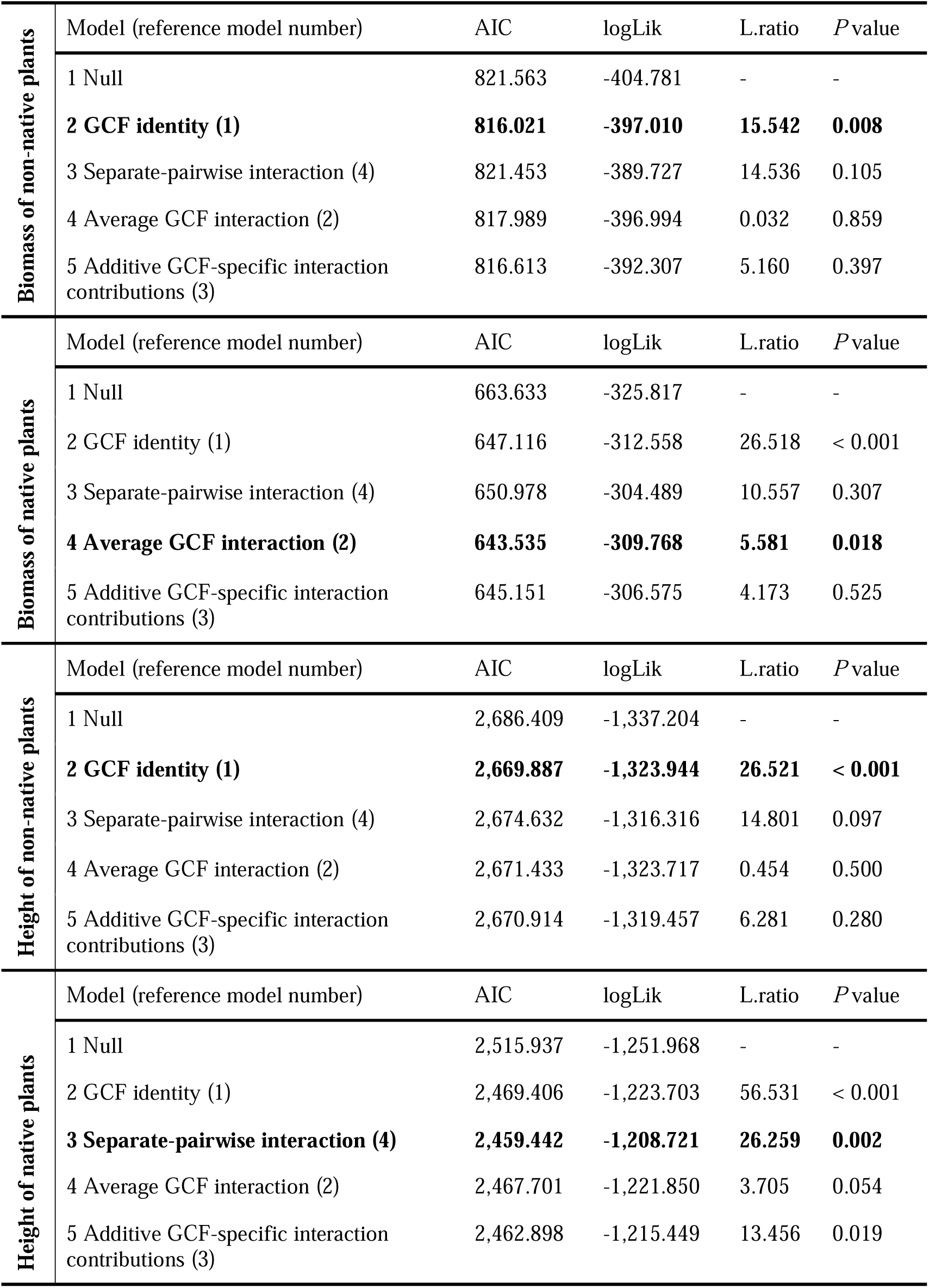
Contributions of GCF identity and GCF interactions to GCF-number effects on the biomass and height of non-native and native plants (see Figure S4 for an explanation of the different models). Model comparisons were based on the log-likelihood-ratio test, and the number in brackets after each model indicates the reference model. The best-fitting model (i.e. with the lowest AIC) is highlighted in bold.

|  |  |  |  |  |  |
| --- | --- | --- | --- | --- | --- |
| Biomass of non-native plants | Model (reference model number) | AIC | logLik | L.ratio | *P* value |
|  | 1 Null | 821.563 | -404.781 | - | - |
|  | **2 GCF identity (1)** | **816.021** | **-397.010** | **15.542** | **0.008** |
|  | 3 Separate-pairwise interaction (4) | 821.453 | -389.727 | 14.536 | 0.105 |
|  | 4 Average GCF interaction (2) | 817.989 | -396.994 | 0.032 | 0.859 |
|  | 5 Additive GCF-specific interaction contributions (3) | 816.613 | -392.307 | 5.160 | 0.397 |
| Biomass of native plants | Model (reference model number) | AIC | logLik | L.ratio | *P* value |
|  | 1 Null | 663.633 | -325.817 | - | - |
|  | 2 GCF identity (1) | 647.116 | -312.558 | 26.518 | < 0.001 |
|  | 3 Separate-pairwise interaction (4) | 650.978 | -304.489 | 10.557 | 0.307 |
|  | **4 Average GCF interaction (2)** | **643.535** | **-309.768** | **5.581** | **0.018** |
|  | 5 Additive GCF-specific interaction contributions (3) | 645.151 | -306.575 | 4.173 | 0.525 |
| Height of non-native plants | Model (reference model number) | AIC | logLik | L.ratio | *P* value |
|  | 1 Null | 2,686.409 | -1,337.204 | - | - |
|  | **2 GCF identity (1)** | **2,669.887** | **-1,323.944** | **26.521** | **< 0.001** |
|  | 3 Separate-pairwise interaction (4) | 2,674.632 | -1,316.316 | 14.801 | 0.097 |
|  | 4 Average GCF interaction (2) | 2,671.433 | -1,323.717 | 0.454 | 0.500 |
|  | 5 Additive GCF-specific interaction contributions (3) | 2,670.914 | -1,319.457 | 6.281 | 0.280 |
| Height of native plants | Model (reference model number) | AIC | logLik | L.ratio | *P* value |
|  | 1 Null | 2,515.937 | -1,251.968 | - | - |
|  | 2 GCF identity (1) | 2,469.406 | -1,223.703 | 56.531 | < 0.001 |
|  | **3 Separate-pairwise interaction (4)** | **2,459.442** | **-1,208.721** | **26.259** | **0.002** |
|  | 4 Average GCF interaction (2) | 2,467.701 | -1,221.850 | 3.705 | 0.054 |
|  | 5 Additive GCF-specific interaction contributions (3) | 2,462.898 | -1,215.449 | 13.456 | 0.019 |

### 3.3 Species-specific growth responses to multiple GCFs

Among the four non-native plant species in the mesocosm community, biomass increased for *Amaranthus retroflexus* (*p* = 0.019) and tended to do so for *Lepidium virginicum* (*p* = 0.081) with increasing GCF number, while there was an opposite trend for *Lolium multiflorum* (*p* = 0.060), and *Senecio inaequidens* showed no significant pattern (Fig. 4A-D). Among the four native plant species, *Silene vulgaris* produced less biomass as the number of GCF increased (*p* = 0.001), while the other three species did not show any significant patterns (Fig. 4I-L). In addition, whereas among the non-native plants only *S. inaequidens* tended to decrease in plant height when the GCF number increased (*p* = 0.071, Fig. 4H), three of the four native species showed a reduction (*Daucus carota*: *p* < 0.001, *S. vulgaris*: *p* = 0.008) or a trend for a reduction (*Plantago lanceolata*: *p* = 0.077) in plant height with increasing GCF number (Fig. 4N-P).

**Figure 4.**
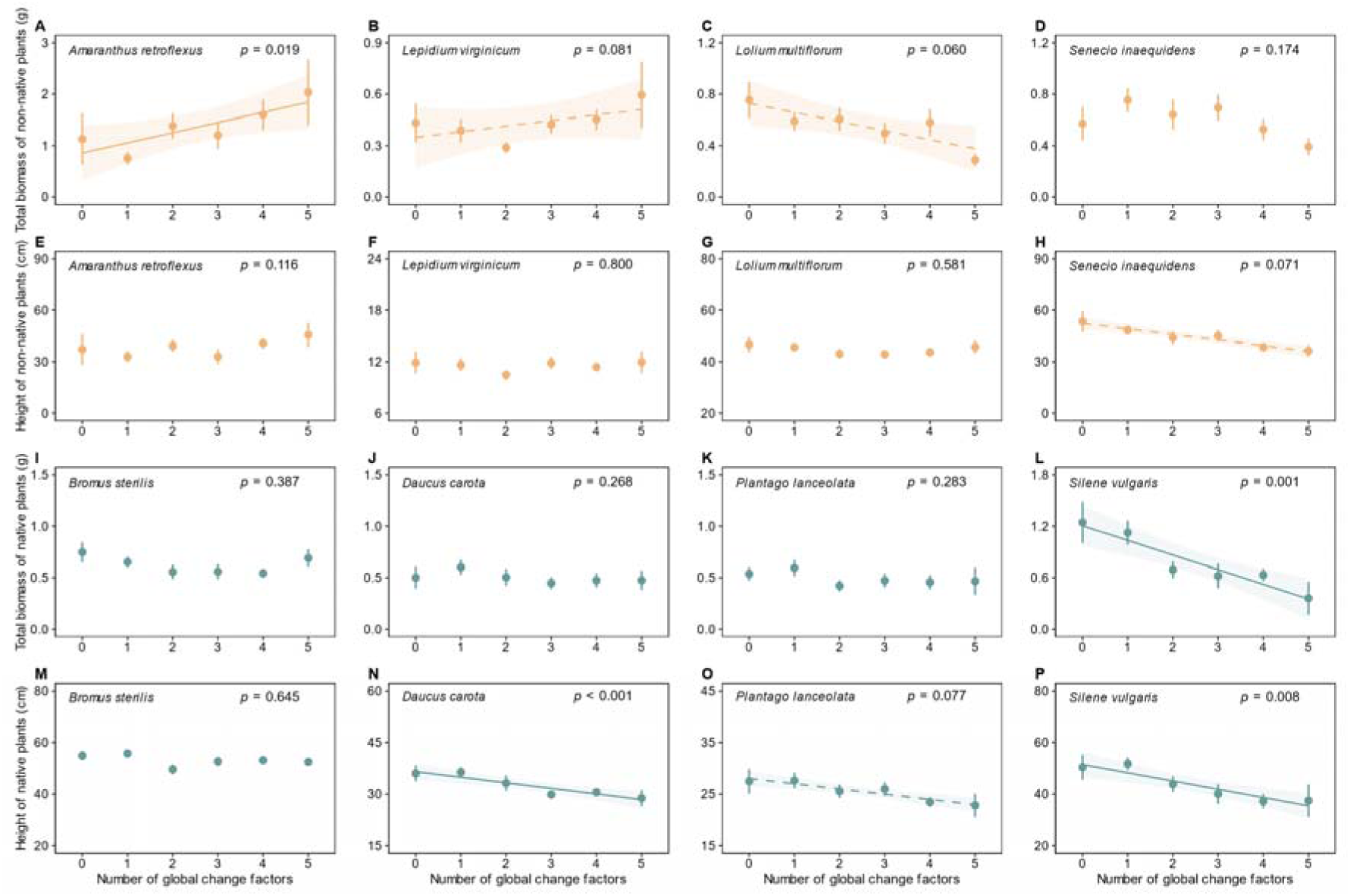
GCF-number effects on biomass and height of the four non-native and native plant species. Data are presented as mean values ± 1 SE. Solid lines indicate significant relationships (*p* < 0.05) with their 95% confidence intervals, and dashed lines indicate marginally significant relationships (0.05 ≤ *p* < 0.1).

## 4. Discussion

Our mesocosm experiment showed that an increase in the number of GCFs had a positive effect on invasion success of the non-native plants. The observed increase in the non-native biomass proportion of the community from c. 42.8% in the control treatment to over 60% in the five-GCF treatment was caused both by an increase in biomass of the non-native plants and a decrease in biomass of the native plants. This is consistent with scenario I in Fig. 1B. For height, on the other hand, we found a negative effect of GCF number for both native and non-native plants, but it was stronger for native plants. This is consistent with scenario V in Fig. 1B. As overall, the native plants were taller than the non-native plants, the difference in height became smaller with increasing GCF number. So, overall, we found that non-native plants a performed better over native plants when the number of GCFs increased.

Although the productivity of our mixed native-non-native community was not significantly affected by GCF number, we found that the dominance of non-native plants in the community was significantly promoted by an increase in GCF number. Hierarchical diversity-interaction models revealed that the GCF-identity model best explained the non-native biomass proportion. This result indicates that the GCF-number effect is mainly due to the additive effects of the single GCFs. Separate analyses for the biomass of non-native plants and native plants showed that the GCF-identity model also best explained the non-native biomass but that the average-GCF-interaction model best explained the native biomass. This suggests that at least for native biomass the negative effect of GCF number is partly due to non-additive effects of GCFs, and that the pairwise interactions were not specific to any GCF. For height of the native plants, there was also evidence for non-additive effects of GCFs, but in this case the contributions of the pairwise interactions depended on the identities of the GCFs. So, although most of the GCF-number effects were due to additive effects of single GCFs, we also found evidence that non-additive effects due to interactions between GCFs may arise, which will make the invasion risk less predictable.

Numerous studies have revealed that non-native plants could benefit from the single GCFs used in our study: nitrogen deposition (Liu *et al*., 2018), ALAN (Liu *et al*., 2022; Kawawa Abonyo & Oduor, 2024), microplastic pollution (Menicagli *et al*., 2023), drought (Carrara *et al*., 2024) and salinization (Bollen *et al*., 2016). To the best of our knowledge, only few studies have tested interactions between two GCFs on plant invasion (Kolb & Alpert, 2003; Lu *et al*., 2022; Yang *et al*., 2022), and no study has previously tested how the number of GCFs influences the performance of non-native plants. Our finding that non-native plants benefit from the number of GCFs, is in line with the findings of many of the studies on single GCFs, but the question that then emerges is why the non-native species benefit while the native species do not. One reason could be that the non-native species have been released from their natural enemies, allowing them to allocate more resources to growth and take more advantage of the GCFs in our study (Xiao *et al*., 2020; Yin *et al*., 2023). On the other hand, recent studies found that both native plant diversity index (evenness and Shannon-Wiener) and resistance of ecosystem service were declined with increasing number of GCFs (Speißer *et al*., 2022; Zhou *et al*., 2024). Consequently, a reduction in community resistance may also increase the possibility of invasion by potential invaders.

Our study found that both biomass and height of the native plants decreased with increasing GCF number. In contrast, Speißer *et al*. (2022) found a positive effect of GCF number on native community productivity. In that study, the effect of GCF number on productivity was mainly driven by a strong positive effect of nitrogen deposition, and the fact that this GCF was included in most of the high GCF-number treatments (Speißer *et al*. 2022). Our study also included nitrogen deposition, but surprisingly it had no significant effect on productivity, possibly because we applied it as a slow-release fertilizer, whereas Speißer *et al*. (2022) applied it as a liquid fertilizer. The negative effect of GCF number on performance of the native plants could be because the GCFs had direct negative effects on the native plants, or because the GCFs had positive effects on the non-native plants resulting in competitive suppression of the native plants (Zhang *et al*., 2021). In line with this, a previous study showed that warming and invasion by *Calluna vulgaris* L. synergistically increased pollinator competition for the native plant *Dracophyllum subulatum* (Giejsztowt *et al*., 2020). Furthermore, it was found that warming and drought in conjunction with invasion by *Pennisetum ciliare* synergistically accelerated the degradation of a native ecosystem (Ravi *et al*., 2022). Therefore, although global environmental changes and plant invasion have been regarded as main drivers of native biodiversity loss (Carboni *et al*., 2021; Pereira *et al*., 2024), the previous studies and our study indicate that multiple GCFs and plant invasion might jointly accelerate this process.

Interestingly, although the biomass proportion of non-native plants increased with the number of GCFs, among the single-GCF treatments only the presence of microplastic had a significant positive effect. Nevertheless, microplastic pollution was not solely responsible for the positive GCF-number effect, because when we excluded all treatments that contained microplastic, the positive effect of GCF number on non-native biomass proportion remained (Fig. S6). The microplastic that we used, EPDM, was previously shown to have negative effects on the growth of native plants, including one of our study species, *D. carota* (van Kleunen *et al*., 2020).

Although we are not aware of any study that tested the effect of EPDM on both native and non-native species growing together, a recent study showed that another non-biodegradable plastic as well as a biodegradable plastic increased the competitive ability of the non-native plant species *Carpobrotus* sp. over the native species *Thinopyrum junceum* (Menicagli *et al*., 2023). The EPDM in our study might have had a direct positive effect on the non-native plants, or, more likely, the non-native plants might have benefited from the negative effect of EPDM on the native plants.

As our experimental community had four non-native and four native plant species, the change in non-native biomass proportion was not necessarily caused by all eight species. Indeed, only one of the native species, *S. vulgaris*, showed a significant negative biomass response to an increase in GCF number. On the other hand, only one of the non-native species, *A. retroflexus*, showed a significant positive biomass response, whereas there were no positive and negative trends for the non-native plants *L. virginicum* and *L. multiflorum*, respectively. However, for the effect of GCF number on height, the responses of the single species were more consistent, as three of the native species (*D. carota*, *S. vulgaris* and *P. lanceolata*) showed significant or marginally significant negative responses, whereas only one non-native species (*S. inaequidens*) showed such a negative trend. Although the multiple GCFs in this experiment did not promote the invasion of all non-native species, three out of four native plants exhibited negative responses in biomass or height. This suggests that multiple GCFs may facilitate the invasion of certain non-native species, which in the long run may lead to the establishment of monocultures of invasive species.

In conclusion, whereas previous studies already showed that the number of GCFs may affect ecosystem service (Zhou *et al*., 2024), soil ecological processes (Rillig *et al*., 2019b; Sáez-Sandino *et al*., 2024), plant individual growth (Zandalinas *et al*., 2021) and native plant community characteristics (Speißer *et al*., 2022), we here show that it might also increase the dominance of non-native plants. However, we also show that the effects of GCF number might vary among non-native species and among native species. Moreover, as mentioned in methods part, our experiment was limited to five global change factors due to constraints in available equipment, and consequently did not include many other potentially important ones. Therefore, future studies should include more non-native and native species and a larger pool of GCFs.

Nevertheless, our results show that the invasion risk of certain non-native species is likely to further increase as the number of global change factors (GCFs) to which plant communities are exposed continues to rise.

## Acknowledgements

We are very grateful to Otmar Ficht, Heinz Vahlenkamp, Kristina Schlötter, Cathrin Wagner, Kyra Boschert and Anna Rankl for their practical assistance. XS acknowledges funding from the China Scholarship Council (202204910008). XS and DC acknowledge the support from the International Max Planck Research School for Quantitative Behaviour, Ecology and Evolution.

## Author contributions

MvK, DC, XS conceived the idea and designed the study, DC and XS performed the experiment and conducted the statistical analysis. XS drafted the first version of the manuscript, with contributions by DC and MvK.

---

**Figure S1.**
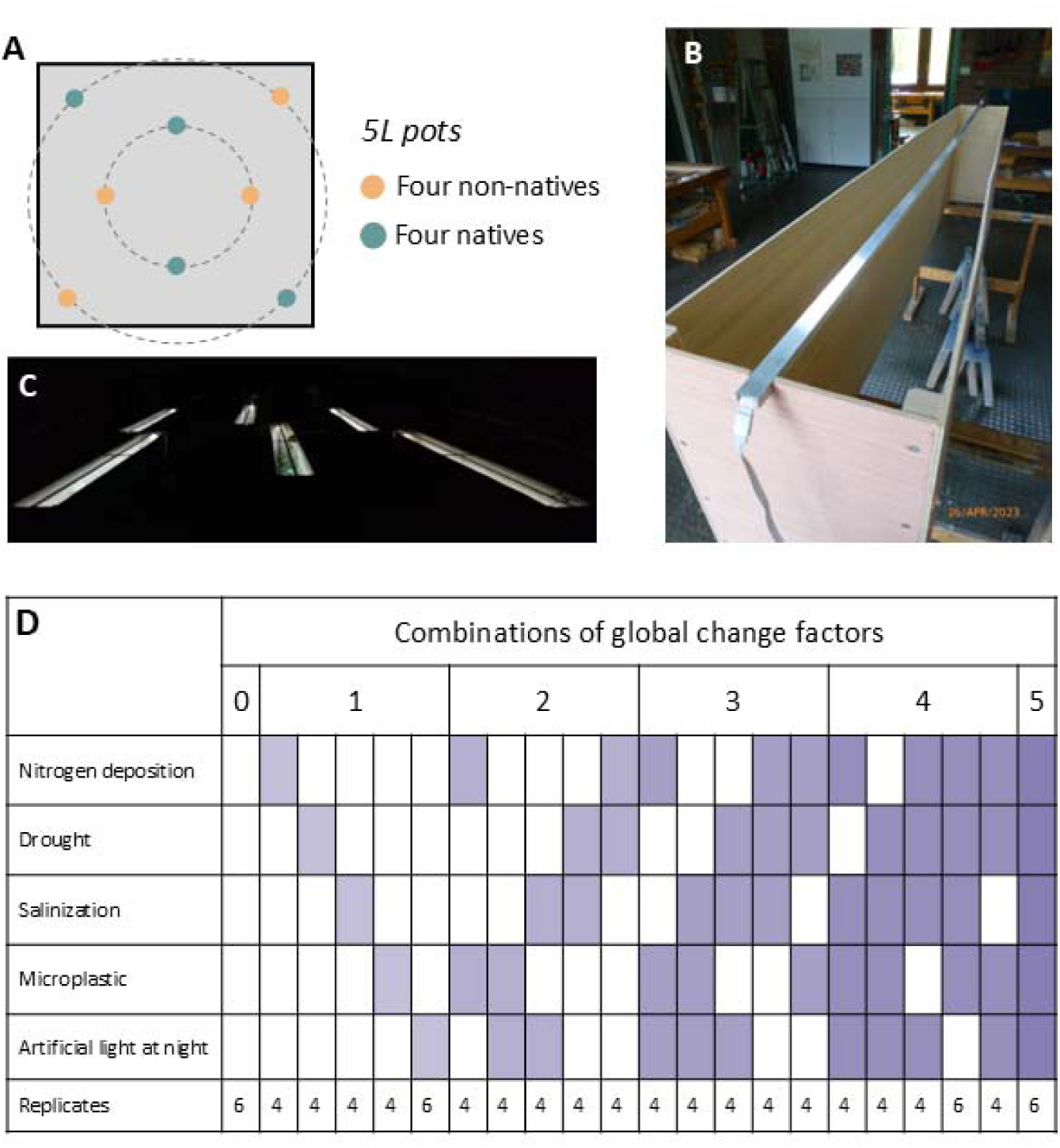
Illustration of the experimental setup. (A), the transplanting positions of the four non-native and four native plants in each pot, they were randomly distributed to four positions for each group. (B), photo of a wooden box with LED strip on the top, C, photo of the experiment at night (next to each lighted box is a control box without light), D, combinations of the five global change factors (GCFs) used to create six GCF-number levels.

**Figure S2.**
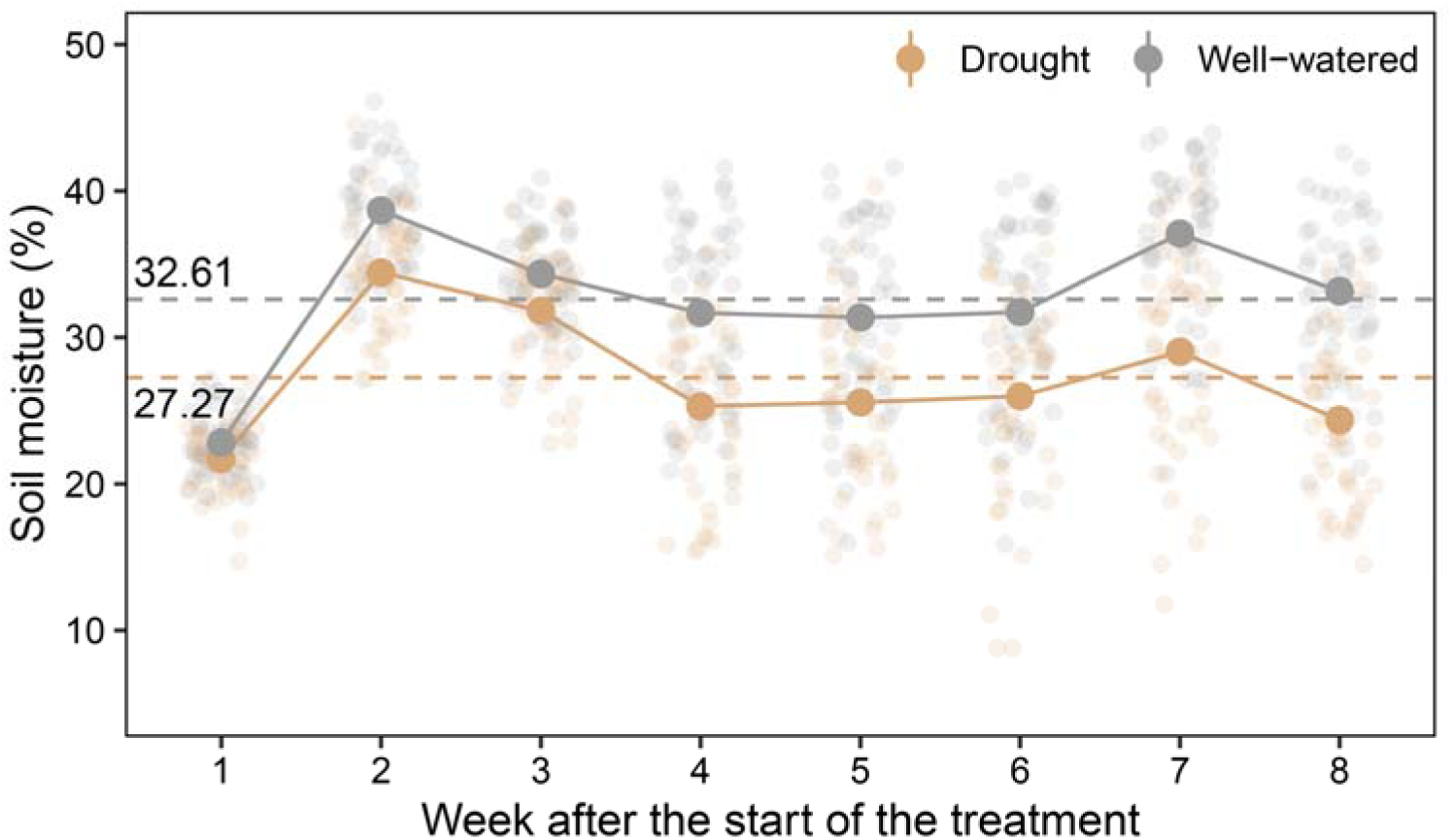
Soil moisture content in the drought and well-watered treatments as measured weekly throughout the experiment. The horizontal dashed lines and the accompanying numbers represent the average soil moisture values for pots in the drought treatment and the well-watered pots across the entire duration of the experiment. Data are presented as mean values ± 1 SE. Smaller circles represent the raw data distribution.

**Figure S3.**
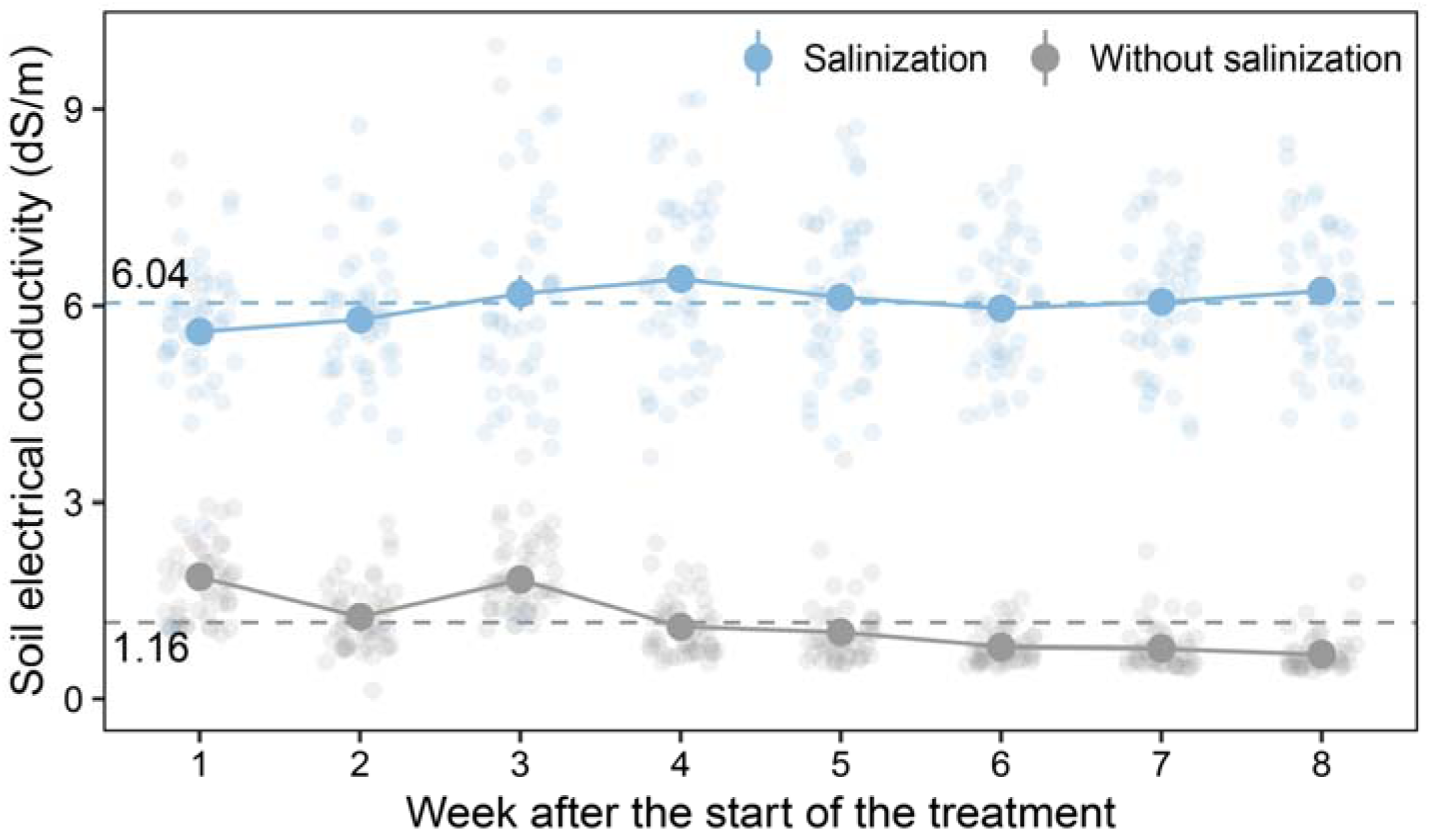
Soil electrical conductivity in the treatments with and without salinization, as measured weekly throughout the experiment. The horizontal dashed lines and the accompanying numbers represent the average soil electrical conductivity values for pots with and without salinization across the entire duration of the experiment. Data are presented as mean values ± 1 SE. Smaller circles represent the raw data distribution.

**Figure S4.**
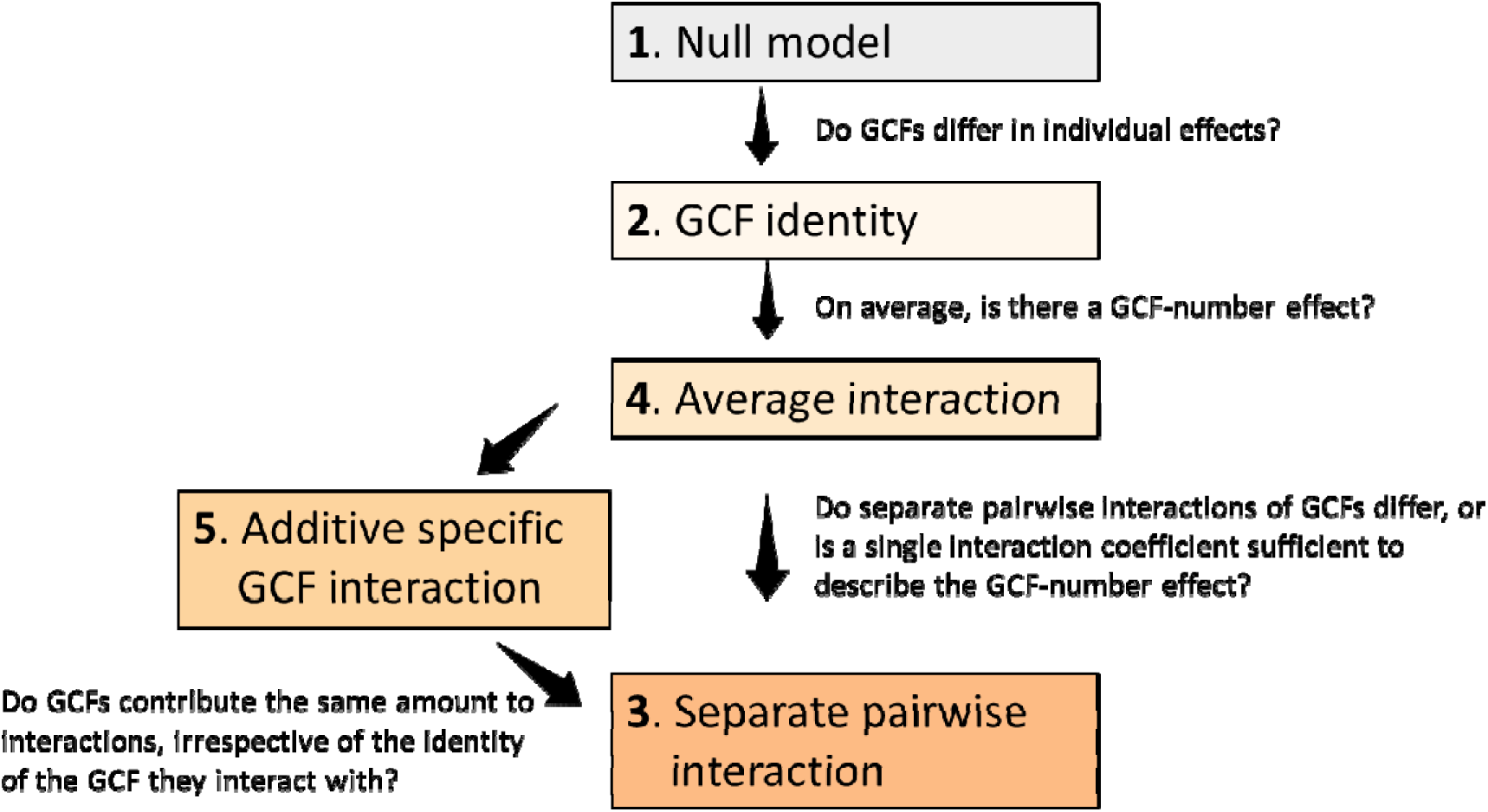
Hierarchical diversity-interaction-modeling framework to assess contributions of GCF identities and GCF interactions to GCF-number effects. The framework, also used by Speißer et al. (2022), is adapted from Kirwan et al. (2009). The null model assumes that there were no interactions between GCFs, and that all GCFs contribute equally. The subsequent models assume more complex effects of how the individual GCFs and their interactions contribute to the GCF-number effects. The interactions can either be specific for each pairwise combination of GCFs (separate pairwise interaction), be the same for each pairwise interaction (average interaction) or be specific for each GCF (additive specific GCF interaction). The questions that can be answered by comparing specific models are indicated by the arrows connecting them.

**Figure S5.**
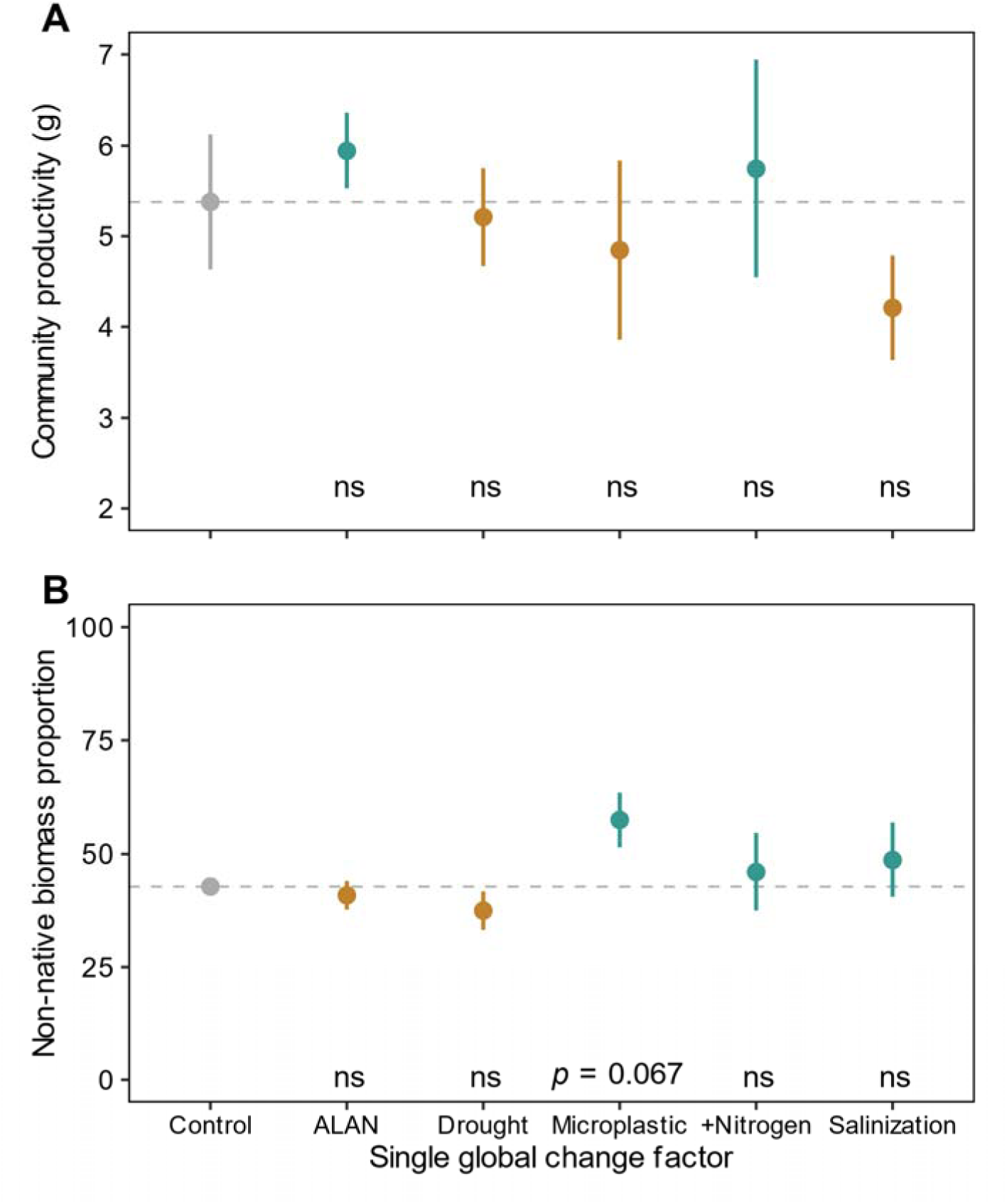
Effects of single GCFs on community productivity (A) and biomass proportion of non-native plants (B). Data are presented as mean values ± 1 SE. Dashed lines and the gray points indicate the mean value in the Control treatment. Green and brown points indicate where the mean value is above or below the dashed line. The *p* values indicate significant differences (*p* < 0.05) between Control and the specific single GCF treatment, ns means no significant difference.

**Figure S6.**
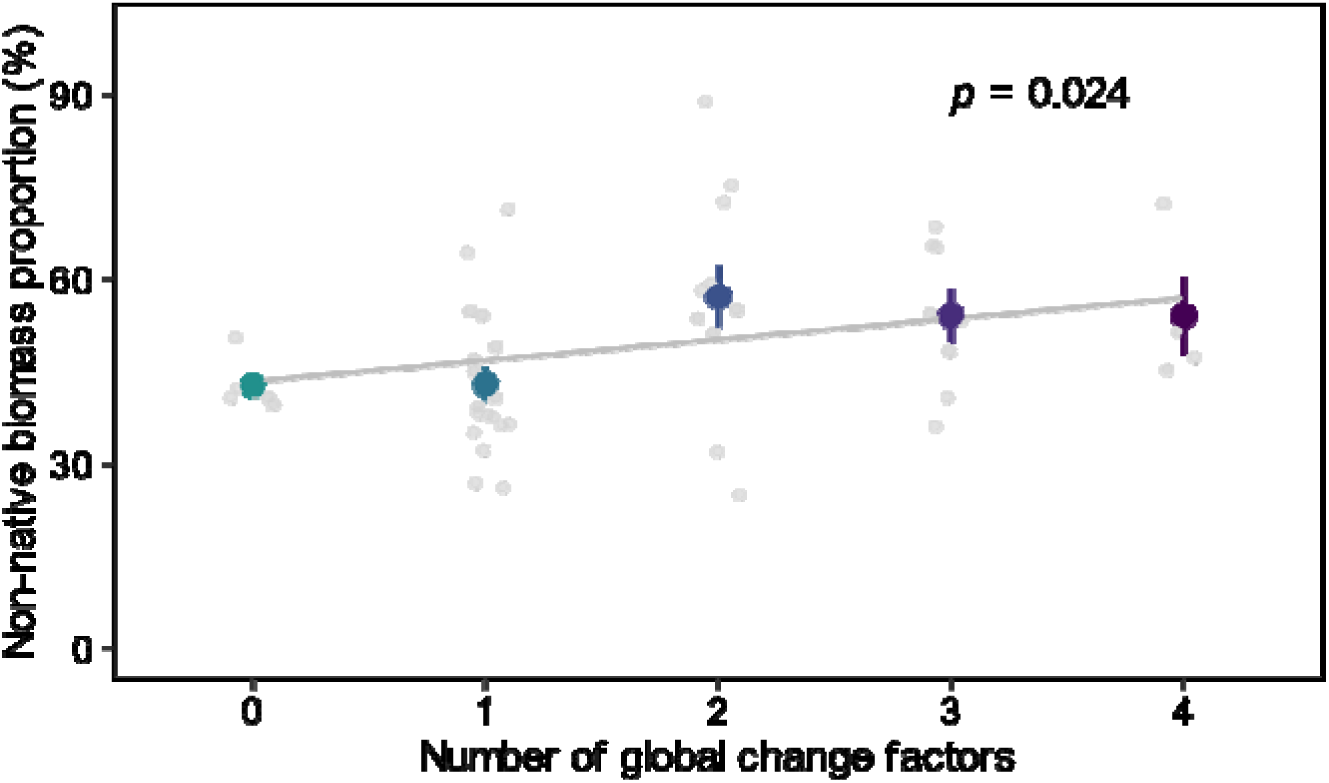
GCF-number effects on biomass proportion of non-native plants. This was done by using the subset of data without the factor microplastics and without combinations that included microplastics. Data are presented as mean values ± 1 SE. Smaller light purple dots indicate the raw data distribution. The line indicates the fitted relationship with its 95% confidence interval. The *p* value indicates the significance of the GCF-number effect.

**Figure S7.**
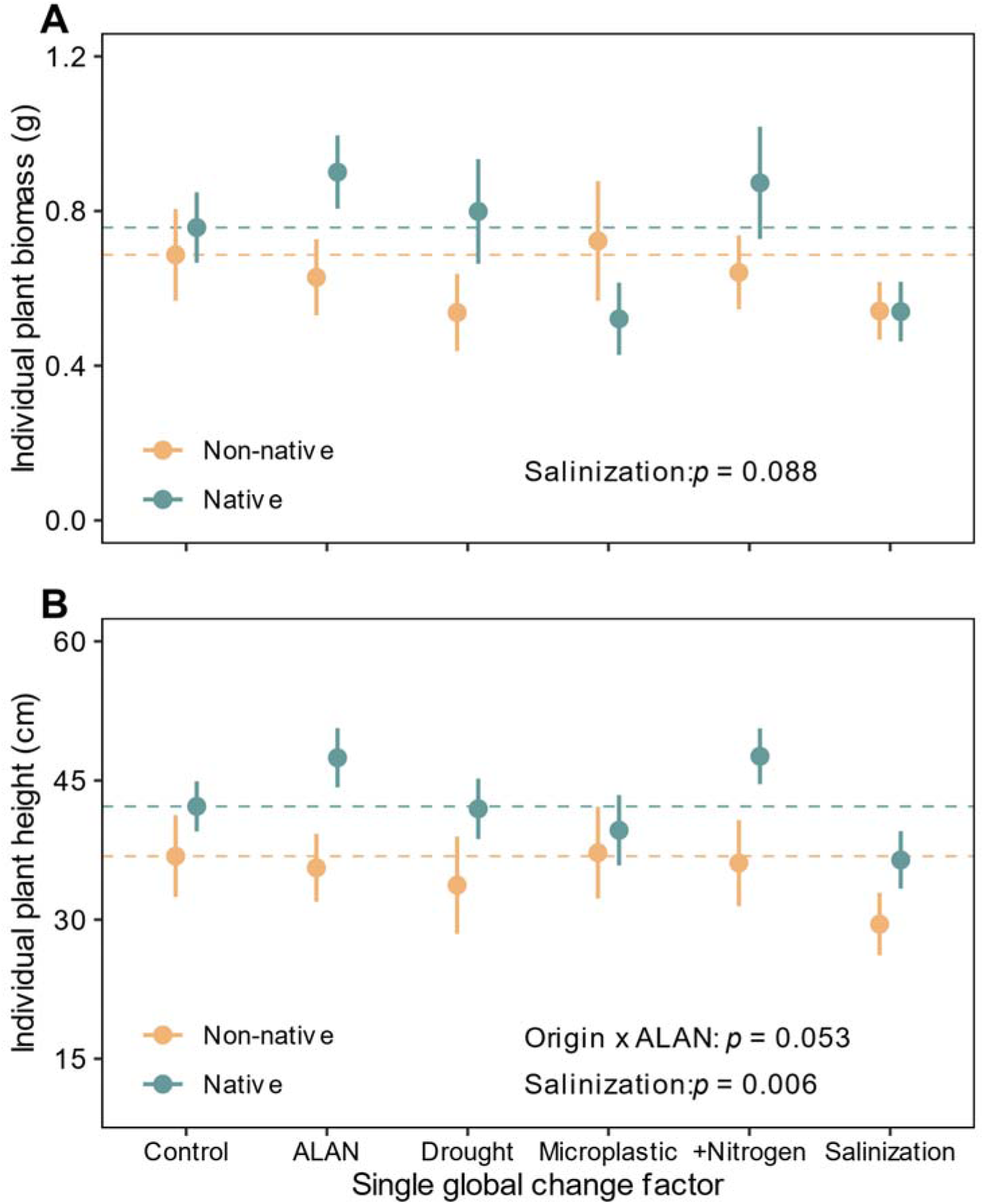
Effects of single GCFs on total biomass (A) and height (B) of non-native and native plants. Data are presented as mean values ± 1 SE. The dashed lines indicate the mean values in the Control treatment. The *p* value indicates significant difference (*p* < 0.05) and marginally significance (0.05 ≤ *p* < 0.1) of individual GCFs and the interaction effects between origin and each GCF based on dummy contrasts.

**Table S1.** Information about the four non-native and four native plant species used in the experiment.

| Species | Family | Origin | Life span | Growth form |
| --- | --- | --- | --- | --- |
| *Amaranthus retroflexus* L. | Amaranthaceae | Non-native | Annual | Forb |
| *Lepidium virginicum* L. | Brassicaceae | Non-native | Annual/Biennial | Forb |
| *Lolium multiflorum* L. | Poaceae | Non-native | Biennial | Grass |
| *Senecio inaequidens* DC. | Asteraceae | Non-native | Perennial | Forb |
| *Bromus sterilis* L. | Poaceae | Native | Annual | Grass |
| *Daucus carota* L. | Apiaceae | Native | Biennial | Forb |
| *Plantago lanceolata* L. | Plantaginaceae | Native | Perennial | Forb |
| *Silene vulgaris* (Moench) Garcke | Caryophyllaceae | Native | Perennial | Forb |

**Table S2.**
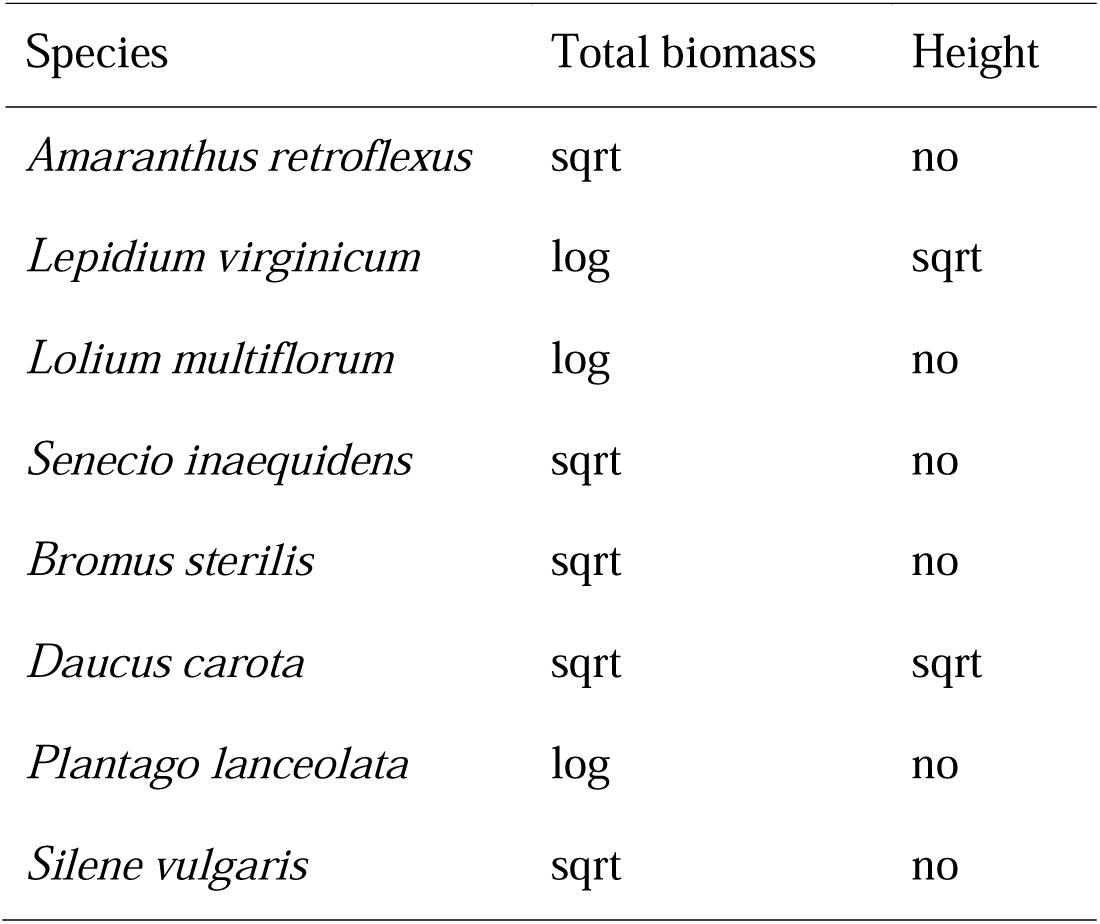
Details about data transformations in the biomass and height models of the eight species (sqrt: square root scale; log: logit scale; no: no transformation).

**Table S3.** Effects of single GCF treatments on community productivity and biomass proportion of non-native plants. Significant effects (*P* < 0.05) are in bold and marginally significant effects (0.05 ≤ *P* < 0.1) are underlined.

|  | Community productivity |  |  | Biomass proportion of non-native plants |  |
| --- | --- | --- | --- | --- | --- |
| *Fixed effects* | *df* | $\chi^2$ | *P* | $\chi^2$ | *P* |
| ALAN | 1 | 0.628 | 0.428 | 0.183 | 0.669 |
| Drought | 1 | 0.065 | 0.798 | 1.103 | 0.294 |
| Microplastics | 1 | 0.500 | 0.480 | 3.365 | <u>0.067</u> |
| Nitrogen deposition | 1 | 0.320 | 0.572 | 0.110 | 0.740 |
| Salinization | 1 | 2.291 | 0.130 | 0.224 | 0.636 |
| *Random effects* |  | *SD* |  | *SD* |  |
| Block |  | 1.026 |  | 0.073 |  |
| Residuals |  | 1.231 |  | 0.229 |  |

**Table S4.** Effects of the GCF number on community productivity and biomass proportion of non-native plants. Significant effects (*P* < 0.05) are in bold.

|  |  | Community productivity |  | Biomass proportion of non-native plants |  |
| --- | --- | --- | --- | --- | --- |
| *Fixed effects* | *df* | $\chi^2$ | *P* | $\chi^2$ | *P* |
| GCF number | 1 | 0.594 | 0.441 | 9.586 | **0.002** |
| *Random effects* |  | *SD* |  | *SD* |  |
| GCF treatment |  | 0.848 |  | 5.489 |  |
| Block |  | 0.616 |  | 4.148 |  |
| Residuals |  | 1.276 |  | 12.757 |  |

**Table S5.** Effects of the GCF number on biomass proportion of non-native plants (this was done in subset of dataset without microplastic treatment and combinations including microplastic). Significant effects (*P* < 0.05) are in bold.

| Biomass proportion of non-native plants |  |  |  |
| --- | --- | --- | --- |
| *Fixed effects* | *df* | $\chi^2$ | *P* |
| GCF number | 1 | 5.109 | **0.024** |
| *Random effects* |  | *SD* |  |
| GCF treatment |  | 5.642 |  |
| Block |  | 5.504 |  |
| Residuals |  | 10.350 |  |

**Table S6.** Effects of species origin (non-native or native), five single GCF treatments and their interactions on plant total biomass and height. Significant effects (*P* < 0.05) are in bold, and marginally significant effects (0.05 ≤ *P* < 0.1) are underlined.

|  | Total biomass |  |  | Height |  |
| --- | --- | --- | --- | --- | --- |
| *Fixed effects* | *df* | $\chi^2$ | *P* | $\chi^2$ | *P* |
| Origin | 1 | 0.672 | 0.412 | 0.508 | 0.476 |
| ALAN (A) | 1 | 0.244 | 0.621 | 0.809 | 0.368 |
| Drought (D) | 1 | 0.416 | 0.519 | 0.298 | 0.585 |
| Microplastics (M) | 1 | 1.661 | 0.197 | 0.270 | 0.603 |
| Nitrogen deposition (N) | 1 | 0.171 | 0.679 | 0.838 | 0.360 |
| Salinization (S) | 1 | 2.902 | <u>0.088</u> | 7.673 | **0.006** |
| Origin × A | 1 | 1.919 | 0.166 | 3.757 | <u>0.053</u> |
| Origin × D | 1 | 1.291 | 0.256 | 0.465 | 0.495 |
| Origin × M | 1 | 1.736 | 0.188 | 0.673 | 0.412 |
| Origin × N | 1 | 0.298 | 0.585 | 2.342 | 0.126 |
| Origin × S | 1 | 0.385 | 0.535 | 0.213 | 0.645 |
| *Random effects* |  | *SD* |  | *SD* |  |
| Species identity |  | 0.118 |  | 15.087 |  |
| Pot identity |  | 0.048 |  | 2.536 |  |
| Block |  | 0.078 |  | 2.239 |  |
| Residuals |  | 0.232 |  | 8.274 |  |

**Table S7.** Effects of species origin (non-native or native), the GCF number and their interaction on plant total biomass and height. Significant effects (*P* < 0.05) are in bold.

|  | Total biomass |  |  | Height |  |
| --- | --- | --- | --- | --- | --- |
| *Fixed effects* | *df* | $\chi^2$ | *P* | $\chi^2$ | *P* |
| Origin | 1 | 0.036 | 0.850 | 0.208 | 0.648 |
| GCF number | 1 | 0.772 | 0.380 | 2.439 | 0.118 |
| Origin × GCF number | 1 | 9.387 | **0.002** | 7.986 | **0.005** |
| *Random effects* |  | *SD* |  | *SD* |  |
| Species identity |  | 0.287 |  | 13.870 |  |
| Pot identity |  | 0.083 |  | 1.979 |  |
| GCF treatment |  | 0.143 |  | 3.552 |  |
| Block |  | 0.170 |  | 2.122 |  |
| Residuals |  | 0.696 |  | 9.182 |  |

**Table S8.**
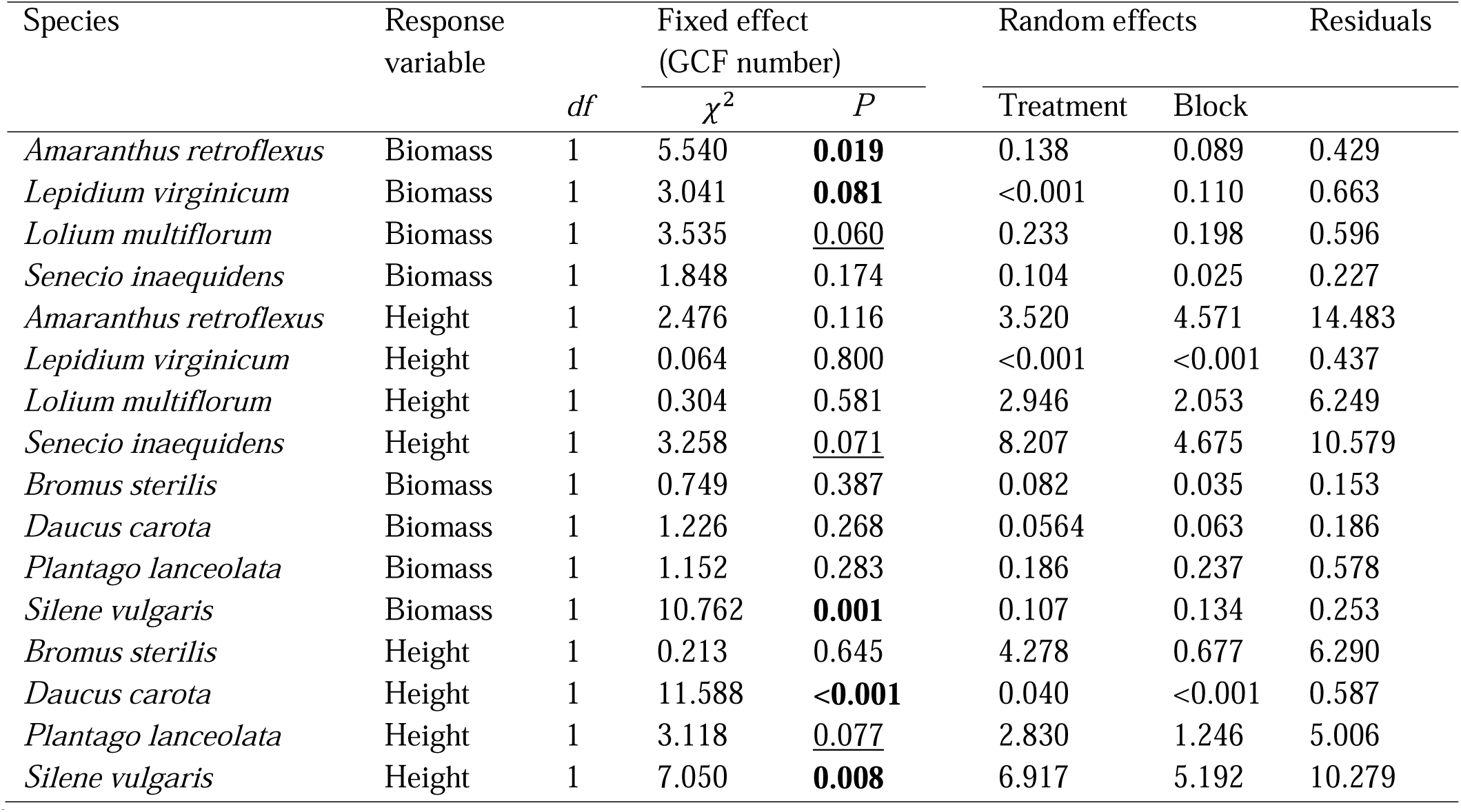
Effects of the GCF number on the total biomass and height of eight plant species (the first four species are non-native, the latters are native). Significant effects (*P* < 0.05) are in bold, and marginally significant effects (0.05 ≤ *P* < 0.1) are underlined.

| Species | Response variable | df | Fixed effect<br>(GCF number) |  | Random effects |  | Residuals |
| --- | --- | --- | --- | --- | --- | --- | --- |
| | | | $\chi^2$ | *P* | Treatment | Block | |
| *Amaranthus retroflexus* | Biomass | 1 | 5.540 | **0.019** | 0.138 | 0.089 | 0.429 |
| *Lepidium virginicum* | Biomass | 1 | 3.041 | **0.081** | <0.001 | 0.110 | 0.663 |
| *Lolium multiflorum* | Biomass | 1 | 3.535 | <u>0.060</u> | 0.233 | 0.198 | 0.596 |
| *Senecio inaequidens* | Biomass | 1 | 1.848 | 0.174 | 0.104 | 0.025 | 0.227 |
| *Amaranthus retroflexus* | Height | 1 | 2.476 | 0.116 | 3.520 | 4.571 | 14.483 |
| *Lepidium virginicum* | Height | 1 | 0.064 | 0.800 | <0.001 | <0.001 | 0.437 |
| *Lolium multiflorum* | Height | 1 | 0.304 | 0.581 | 2.946 | 2.053 | 6.249 |
| *Senecio inaequidens* | Height | 1 | 3.258 | <u>0.071</u> | 8.207 | 4.675 | 10.579 |
| *Bromus sterilis* | Biomass | 1 | 0.749 | 0.387 | 0.082 | 0.035 | 0.153 |
| *Daucus carota* | Biomass | 1 | 1.226 | 0.268 | 0.0564 | 0.063 | 0.186 |
| *Plantago lanceolata* | Biomass | 1 | 1.152 | 0.283 | 0.186 | 0.237 | 0.578 |
| *Silene vulgaris* | Biomass | 1 | 10.762 | **0.001** | 0.107 | 0.134 | 0.253 |
| *Bromus sterilis* | Height | 1 | 0.213 | 0.645 | 4.278 | 0.677 | 6.290 |
| *Daucus carota* | Height | 1 | 11.588 | **<0.001** | 0.040 | <0.001 | 0.587 |
| *Plantago lanceolata* | Height | 1 | 3.118 | <u>0.077</u> | 2.830 | 1.246 | 5.006 |
| *Silene vulgaris* | Height | 1 | 7.050 | **0.008** | 6.917 | 5.192 | 10.279 |

